# Entangled and non-modular enhancer sequences producing independent spatial activities

**DOI:** 10.1101/2024.07.08.602541

**Authors:** Mariam Museridze, Stefano Ceolin, Bettina Mühling, Srishti Ramanathan, Olga Barmina, Pallavi Santhi Sekhar, Nicolas Gompel

**Affiliations:** Ludwig-Maximilians Universität München, Fakultät für Biologie, Biozentrum, Planegg-Martinsried, Germany; University of Bonn, Bonn Institute for Organismic Biology, Bonn, Germany; Department of Evolution and Ecology, University of California, Davis, California, United States of America

## Abstract

The modularity of transcriptional enhancers (*1, 2*) is central to our understanding of morphological evolution, as it allows specific changes to a gene expression pattern component, without affecting others (*3*). Enhancer modularity refers to physically separated stretches of regulatory sequence producing discrete spatiotemporal transcriptional activity. This concept stems mainly from assays that test the sufficiency of a DNA segment (*4*) to drive spatial reporter expression resembling that of the corresponding gene. Focusing on sufficiency to produce spatial patterns, it overlooks quantitative aspects of gene expression, underestimating the regulatory sequence actually required to reach full endogenous expression levels. Here we show that five regulatory activities of the pigmentation gene *yellow* in *Drosophila*, classically described as modular (*5–7*), result from extensively overlapping sequences, with broadly distributed regulatory information. Nevertheless, the independent regulatory activities of these entangled enhancers appear to be nucleated by specific segments that we called enhancer cores. Our work calls for a reappraisal of enhancer definition and properties, as well as of the consequences on regulatory evolution.

## Introduction

The embryonic blueprint of gene expression prefiguring morphology results from the activity of separate *cis*-regulatory elements (*8–12*). It was hypothesized (*13, 14*), then demonstrated (*5, 15–17*) (for review, see (*3*)) that mutations in enhancers are a primary driver for the evolution of discrete morphological traits, such as gains and losses of limbs in vertebrates or decorative elements in insects. Owing to their reduced pleiotropy, mutations affecting single enhancers facilitate changes in specific aspects of morphology with no deleterious effect on other traits (*3, 18*). Hence, a modular representation of enhancers bears strongly on our understanding of regulatory evolution.

The original notion of enhancer modularity (*2*) was quickly generalized in transgenic animals and plants with reporter assays testing the capacity of arbitrarily chosen DNA segments to recapitulate elements of the gene’s spatial expression. This notion is based on the sufficiency of DNA segments to drive specific expression patterns, leaving out completely how much transcript these sequences yield compared to the endogenous levels of transcript. Yet, the amount of gene transcript is key to normal development (*19, 20*). Moreover, a minimal enhancer producing a correct spatio-temporal pattern may be insufficient for a robust expression under unfavorable environmental or genetic conditions (*21–23*). In fact, it has been repeatedly shown that mutations occurring outside minimal enhancers can contribute to phenotypic evolution (*23–25*).

The concept of enhancer modularity was nevertheless further comforted by the advent of genomics, identifying peaks of various epigenetic marks or accessibility in phase with previously identified regulatory segments (*26*). Yet, the extent of sequence that these peaks span is influenced by the choice of cut-off values or other methodological biases. For instance, the processing of datasets assessing genome-wide chromatin accessibility starts with the binning of raw data into putative regulatory elements (*27*), leading to a circular argument. As a result, of these different approaches, transgenic assays or genomic methods, enhancers are deemed to span 100-1000 base pairs, although the sequence necessary for a full regulatory activity –pattern and levels– may be larger (*28*).

Does the picture of discrete and modular enhancers hold when one considers the quantitative dimension of transcription? Using a quantitative reporter assay, we addressed this question by precisely mapping regulatory information at the *yellow* (*y*) locus in *Drosophila*, a typical locus with several independent enhancers. Their activities, classically mapped with reporter assays, prefigure a gray background or a black spot on the wing, stripes, a longitudinal band, a sexually dimorphic block of pigmentation on the abdomen, or black sensory bristles over the body, respectively (*5, 7, 29*). A recent systematic dissection of *y* regulatory regions in *Drosophila melanogaster* (*30*), however, suggested more distributed activities than the textbook picture of separate and independent enhancers would predict. Along the same lines, focusing on the wing-spotted species *Drosophila biarmipes*, we showed that the *wing blade* and *spot* enhancers, respectively prefiguring the background gray pigmentation of the wing and a dark distal wing spot, were actually broader, extensively overlapping, and shared regulatory information (*31*). The sequence overlap of these two enhancers may, however, simply result from a recent cooption process, and the regulatory activities may resolve with enough evolutionary time.

To investigate the relationship between distinct regulatory activities, we compared the wing enhancers to an ancient regulatory sequence, the *body* enhancer. The *body* enhancer, found in various *Drosophila* species (*6, 32*), is active in the head, thoracic and abdominal epidermis during metamorphosis (*33, 34*) defining a complex spatial pattern. We mapped regulatory information of the *body* enhancer and compared it to our previous map of the wing enhancers (*31*). The *body* enhancer spans the entire sequence of the two wing enhancers, refuting their modularity. The sequences of these activities are entangled with, however different distributions of regulatory information.

## Results

*yellow* is expressed in body pupal epidermis (*6, 35*) under the control of regulatory sequences mapping 5’ of *yellow* transcription start site (TSS) (*29, 33, 34*). The dissection of these sequences from *D. melanogaster* with transgenic reporter assays identified a 1 kb segment, the *body* enhancer, sufficient to recapitulate *y* spatial expression (*6*). Whether this segment is sufficient to drive endogenous transcription levels is, however, unknown. We therefore first mapped the entire regulatory sequences necessary and sufficient to drive *yellow* transcription in the fly body, relying on a quantitative reporter assay. For the sake of comparison with our previous mapping, we used *D. biarmipes* sequences, and for simplicity, we focused on abdominal expression. Starting from a 5.4 kb segment upstream of *yellow* transcription start site, we tested the regulatory activity of segments with deletions from the 5’ end (D series), as well as progressive sequence randomization from the 3’ end in male pupae (E series; Fig. 1A, C and ref. (*31*)). To map the boundaries of regulatory activities in the abdomen, we calculated how much reporter expression was lost or gained for each construct compared to the activity of the largest construct of the corresponding series (Fig. 1B, see methods). For instance, removing a distal segment (line *D1*), about 4 kb upstream of the original *body* enhancer (*6*), already had a significant effect on the abdominal expression (Fig. 1B). Generally, we found that all constructs differed in activity from the largest construct of each series (Table S1). We concluded that the regulatory information determining abdominal expression spans up to 5.4 kb and overlaps largely with the previously mapped regulatory activities of the wing (*31*), challenging the notion of short and modular enhancers. Importantly, average phenotypes (Fig. 1A) revealed changes not only in the levels of regulatory activity among constructs, but also in spatial distribution. We therefore sought to understand how the distributed regulatory information upstream of *yellow* TSS was organized to control different spatial pattern elements. To this end, we relied on a principal component analysis (PCA) of phenotypic variation across all constructs (Fig. 2A, B; Fig. S1), as it gave us access to the fine relationship between regulatory content and enhancer activity. The first component (PC1, 72% of the total variation, Fig. 2A, B), mostly captured broad uniform expression changes across the abdomen. The second component (PC2, 18% of the total variation) captured changes in two distinct spatial activities simultaneously, in the upper four segments and the last two segments, respectively corresponding to broad uniform expression (*30*) and the sexually dimorphic male expression (Fig. 2A, B). The third component showed an identified artifact caused by gaps between the last abdominal segments, due to ventral bending (Fig. S1). We did not consider PC3 for further analysis. The fourth component (PC4, 1% of the total variation), shows changes in the basal part of each segment, identifying the banded pattern of the fly abdomen (Fig. 2A). In summary, PC1, PC2 and PC4 appear to jointly capture variation in the three main pattern components of the global *body* activity, although these PCs are not necessarily independent.

**Fig. 1.**
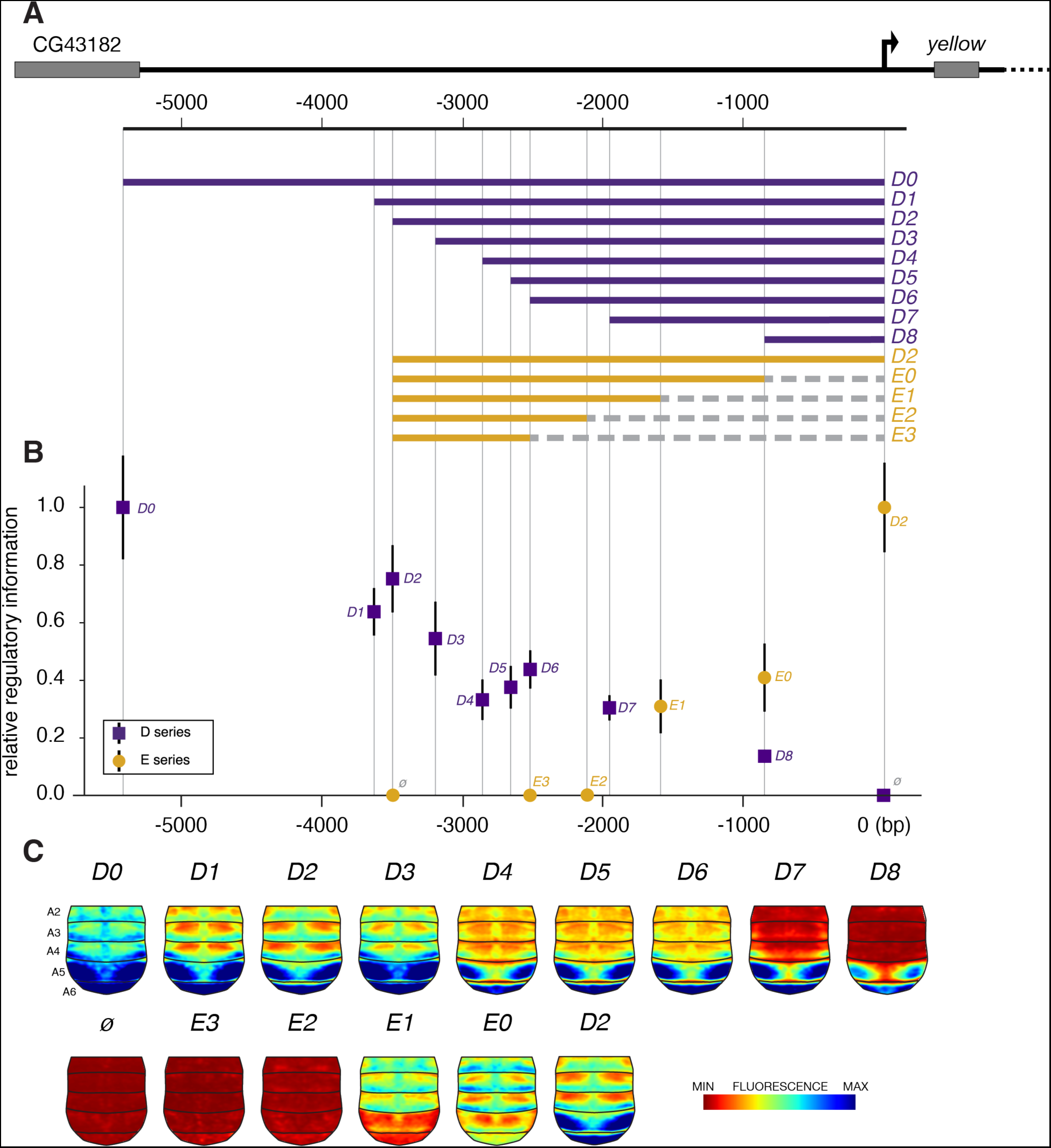
Mapping regulatory activities upstream of the *yellow* gene. (**A**) Reporter construct series used to evaluate the regulatory content of regions located 5’ to *yellow*. A first series, D (purple), is progressively trimmed from the 5’ end, while the proximal end of constructs of the second series, E (ochre), is increasingly randomized, without changing the distance to the transcription start site. Note that construct *D2* appears in the E series, too, as the reference construct from which all E constructs were derived. (**B**) Relative regulatory information (fluorescence levels) for each reporter line compared to the reference line of the series. (**C**) Average reporter expression (average phenotype) in the abdominal epidermis for each line. *ø* denotes a line with an empty reporter vector. In all figures, only male abdomens were examined.

**Fig. 2.**
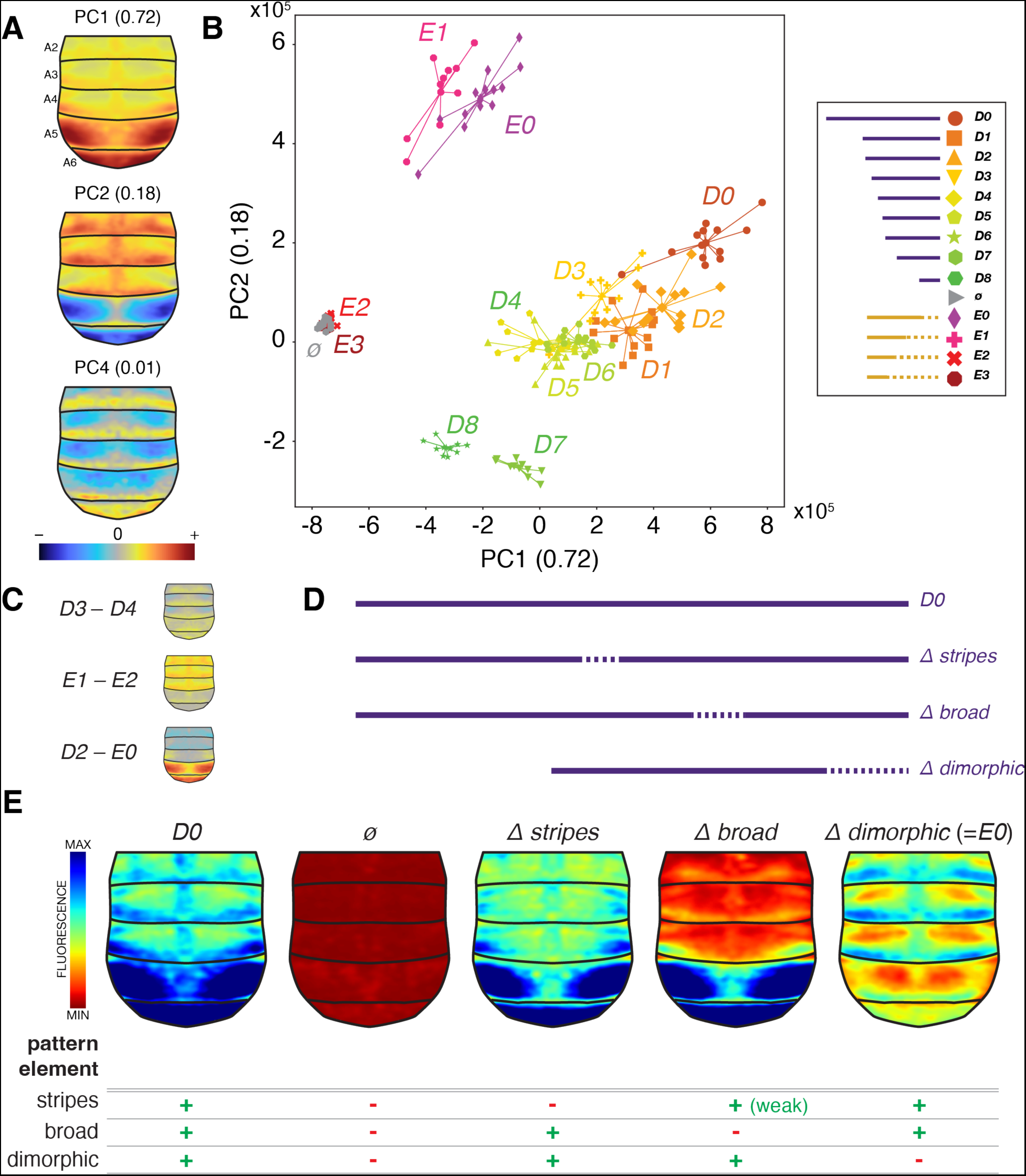
The *body* enhancer pattern results from three distinct activities. (**A**) Phenotypic directions captured by three principal components in the space of variation for all reporter lines shown in Fig. 1. These directions correspond to broad uniform expression (PC1), a combination of broad uniform expression and sexually dimorphic male expression (PC2), and stripes (PC4). PC3 identifies an artifact caused by stretching between consecutive segments and is shown in Fig. S1. (**B**) PCA of phenotypic space showing the first two principal components. Each dot represents a single abdomen. All samples from the same construct are connected by lines to their average. (**C**) Subtraction of average phenotypes of selected lines identifies specific pattern elements: *D3* – *D4*, stripes; *E1 – E2*, broad uniform expression; *D2* – *E0*, sexually dimorphic element. (**D**) Constructs with randomized sequence segments according to (C). (**E**) Resulting average phenotypes of lines with randomized segments with an annotation of pattern elements.

To test the possible regulatory independence of these activities, we sought to identify sequence segments that affect single spatial pattern components. To this end, we first calculated differences between average phenotypes of consecutive lines (Fig. S2). We observed that the average phenotype difference between certain lines corresponded precisely to pure individual activities (Fig. 2C): *D3* – *D4* reflects the stripes component, *E1* – *E2* the broad uniform component, and *D2 – E0* the dimorphic component, suggesting that the corresponding sequence segments may affect the respective activities independently. To directly test these independent regulatory contributions, we examined reporter constructs with randomized sequence in place of these segments (Fig. 2D): *Δ stripes* (segment *D3*-*D4* randomized in the context of *D0*), *Δ broad* (segment *E1-E2* randomized in the context of *D0*), and *Δ dimorphic* (segment *D2-E0* randomized in the context of *D2* = construct *E0* of Fig. 1). Compared to the activity of the largest construct *D0*, which contains all three pattern components, stripes, broad expression, and dimorphic expression, we observed that only a single component was missing in each randomized line (Fig. 2E): *Δ stripes* had no visible stripes, *Δ broad* missed broad expression in the upper segments, and *Δ dimorphic* was devoid of strong expression in the posterior segments. These results showed that the three *body* activities are to some extent functionally independent of each other. Further, they suggested that each of these segments could be an enhancer core, that is a stretch of sequence necessary to seed the spatial regulatory activity, but insufficient to account for the endogenous levels of expression (*31*). To evaluate the extent of independence of the three activities, *broad*, *dimorphic* and *stripes*, and to finally examine the distribution of regulatory information upstream of *yellow* TSS, we went on to decompose the expression patterns as a sum of these three activities in the PC space. The decomposition is achieved through a simple operation, a change of base, whereby the PC space is reprojected in a new coordinate system made of three vectors corresponding to the pure activities. The *stripes* activity is defined by the vector *(D0 to Δ stripes)* in the original PC space, the *broad* activity by the vector *(D0 to Δ broad)*, and the *dimorphic* activity by the vector *(D2 to Δ dimorphic)*. The resulting mixed components of the re-projected space show changes only in stripes, broad, or dimorphic pattern elements, respectively (Fig. 3). In this reprojected space we could then measure the independent contribution of each fragment in the dissection series to each pattern component and compare these distributions of regulatory information to those of the wing enhancers that we calculated in ref. (*31*) (Fig. 3). The results show that, in spite of their extensive overlap, the distribution of regulatory information along the sequence is very different among activities. Together, our results outline an alternative architecture of *y* regulatory regions, whereby enhancer cores necessary to initiate a regulatory activity are surrounded by regulatory sequences contributing to the full endogenous levels of transcription. It is at odds with the current thread-like image of modular regulatory elements at *y*, or generally at developmental loci. This could be a peculiarity of the *y* locus. Yet, the methodological biases of ascertainment caused by classical reporter assays and genomic data processing (*27*) speak for an entangled architecture of regulatory regions in general. This is only comforted by circumstantial evidence that regulatory regions have been underestimated (*23–25, 28*). This emerging possibility raises two fundamental questions.

**Fig. 3.**
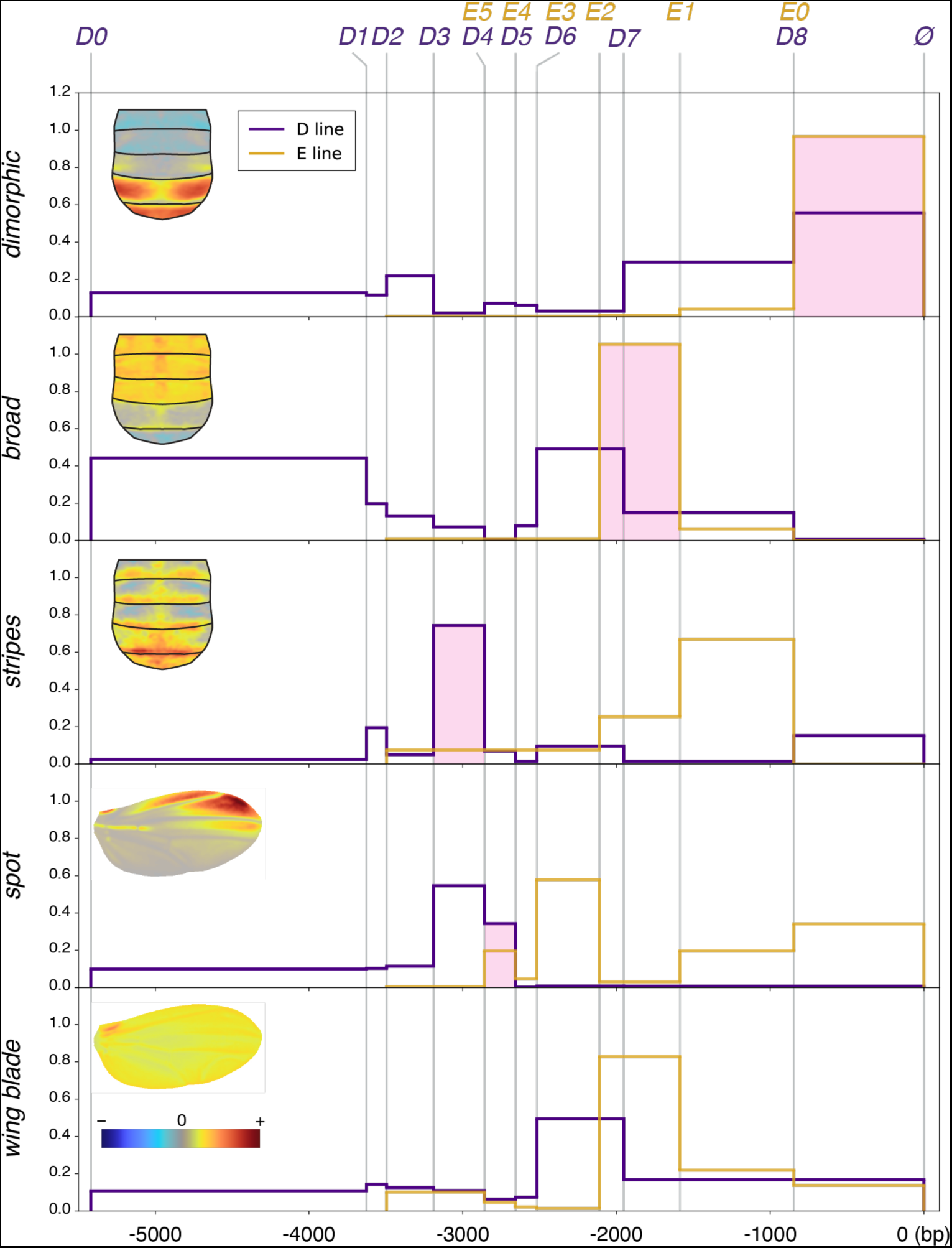
Regulatory architecture upstream of *yellow*. The abdomen and wing images with a colormap correspond to the mixed components, that is the variation of different regulatory activities in the PC space, after a change of base. While principal components of the original PCA captured multiple activities, the mixed components resulting from a change of base capture variation in single activities (*i.e.*, phenotypic directions). The contribution of each sequence segment along *y* 5’ region to each mixed PC is a map of the regulatory information for each enhancer. While all activities span most of the 5.4 kb region, we note that enhancers cores (pink shading) identified in Fig. 2 and ref. (*31*)) are not overlapping.

First, if enhancers are not modular stretches of sequence, then how do they independently produce regulatory activities in distinct subsets of cells? At the *y* locus, each enhancer responds to a subset of transcription factors (TFs) governing its activity, but TFs present in the wing may also be present in the abdomen and could activate wing enhancers in abdominal cells. We compared the TFs expressed in pupal wings (*36, 37*) to those expressed in abdominal epidermis at the same stage (using a previously unpublished micro-array dataset, see methods). Of 302 wing TFs and 240 abdominal TFs, we found that 78 are expressed in both tissues during pupal life (Table S2), potentially exposing their respective enhancers to bleed-through activities.

We submit that the regulatory independence may reside in the function of enhancer cores and hypothesize that each core subordinates all flanking regulatory information through an unknown mechanism to selectively produce the full activity of a given enhancer. This could involve selective accessibility or selective use of transcription factor binding sites, or both, as we have shown in the case of the *spot* enhancer (*31*). With the advent of methodologies describing the dynamics of the 3D genome, the representation of gene regulation is changing from selective and transient looping of a remote enhancer onto the core promoter region, to more stable multi-enhancer hubs gathered around the TSS of a gene (*38, 39*). It is conceivable that a complex architecture of entangled enhancers forms a relatively stable hub around a core promoter, where core enhancer segments control the selective involvement of each activity in the respective cell types.

Second, if enhancers share extensive portions of sequence, how are their independent changes possible, a cornerstone concept of morphological evolution? One possibility is that binding sites for TFs involved in distinct activities are intermingled rather than shared. This would still afford changes in selected activities with minimal effects on other enhancers. On the other hand, if the 3D folding of a regulatory region with entangled enhancers is relevant for its function, the evolvability of regulatory sequences may be more limited than previously assumed.

## Acknowledgments

We are grateful to Benjamin Prud’homme for insightful comments on the manuscript.

## Author contributions

Conceptualization: MM, NG

Methodology: MM, SC

Investigation: MM, SC, BM, SR, PSS, OB

Visualization: MM, NG

Funding acquisition: NG

Project administration: NG

Supervision: NG

Writing – original draft: NG, MM

Writing – review & editing: MM, NG, SC

## Data and materials availability

All data are available in the main text or the supplementary materials.

## Supplementary Materials

### Materials and Methods

#### Fly husbandry

Our *D. melanogaster* stocks were maintained on standard cornmeal medium at 25°C with a 12:12 day:night light cycle.

#### Transgenesis

All reporter constructs were injected as in Arnoult et al. (*36*). We used ɸC31-mediated transgenesis (*40*) and integrated all constructs at the genomic attP site VK00016 on chromosome 2 (*41*). The enhancer sequence of all transgenic stocks was genotyped before imaging.

#### Molecular Biology

Lines from the D and E series and the randomized sequence for the *Δ stripes* and *Δ broad* were taken from (*31*). Fragments of *y* 5’ sequences for lines *D8*, *Δ stripes*, *Δ broad* were amplified with Phusion polymerase (NEB) and cloned into our transformation vector pRedSA (*31*) digested with EcoRI and BamHI using In-Fusion HD Cloning Kits (Takara; catalog no. 121416). Randomized sequences were amplified from the fragment used in (*31*). Primers are listed in Table S3. The sequences are provided in Table S4.

#### Imaging

##### Sample preparation

All transgenic abdomens imaged in this study were from male flies heterozygous for the reporter construct, obtained from a cross between homozygous reporter construct flies and a marker line (;*en*-Gal4, UAS-GFP/CyO; *pnr*-Gal4/TM6B, (*42*), FlyBaseID FBrf0098595). White pupae were left to age 90 h at 25°C. Pupal case was removed and pupae were mounted in halocarbon oil (Sigma-A, CAS-No 9002-83-9) on a microscope slide with cover slips and appropriate spacers, and immediately imaged.

##### Microscopy

All abdomen images were acquired as 12-bit images on a Leica SP5-2 confocal microscope using an HC PL APO CS 10x/0.40 lens with a HyD SP Hybrid-Detector. Each image comprises two fluorescent channels (DsRed to report *yellow* regulatory activities and *en-*Gal4, *pnr*-Gal4, UAS-GFP as positional landmark for image registration). Image stacks were acquired in 4.99 µm steps.

##### Image Quantification and Analysis

We utilized a comprehensive pipeline for the registration, segmentation, and analysis of multi-channel 3D fluorescence microscopy images. This pipeline, currently supporting images with three color channels, facilitates the analysis of gene expression patterns in the epithelial cells of the *Drosophila* abdomen by producing a registered 2D map of expression on the abdomen surface starting from a 3D image stack (Fig. S3A). The pipeline begins with an automated preprocessing step (Fig. S3B), where segmentation of the image stack is performed using an automatically determined threshold. This threshold is set to ensure that the segmented volume covers a specified range of the entire image volume. Post-segmentation, the object is refined using morphological transformations and mesh fitting to fill any holes. Following preprocessing, the registration step involves rendering the segmented volume in an interactive interface. Corresponding points on the source and reference objects are selected, and the image stack is rigidly rotated and rescaled to minimize the distances between these points (Fig. S3C). In the projection phase, the surface of the registered objects is projected into 2D images using a modified sinusoidal projection. The object is sliced, and for each slice, the profile of the bright object is interpolated with a spline curve. Image brightness is then read out along the curve by taking the local maxima along the local normal direction. The resulting 1D brightness profiles from each slice form the rows of the 2D projected image, which are aligned at a predefined meridian plane (Fig. S3D). The final step involves labeling and elastic warping. A graphical interface built with the PYSimpleGUI library enables manual labeling and registration of the 2D images. User-selected points are utilized to elastically warp the images onto a reference model using thin-plate spline registration (Fig. S3E). This methodology ensures precise registration and analysis of multi-channel 3-dimensional images, thereby facilitating the detailed study of gene expression patterns in the *Drosophila* abdomen. The python-based code, together with an example jupyter notebooks, installation instructions and a minimal test dataset is available at https://github.com/UniBonn-GompelLab/3D-DrosophilaRegistration.

##### Relative regulatory information

We showed how much regulatory information (fluorescence levels) is contained in each line and compared it to the construct containing the largest regulatory fragment. First, to measure the amount of fluorescence we calculate a squared Euclidean distance from each sample to the average phenotype of the line with an empty vector (*ø*). Practically, we subtracted from each projected abdomen image the average image of the empty line (*ø*), then squared and sum all the values. Next, we calculated an average value of these distances for each line. Finally, to show the relative amount of regulatory information, we divided all the line averages by the average of the biggest construct in the series. (*D0* line for the D series and *D2* line for the E series). The statistical significance of differences among lines was assessed with Kolmogorov-Smirnov tests (Table S1).

##### Density of regulatory information per base

The amount of regulatory information brought by a segment of DNA for a given activity was calculated as the absolute value of the difference between the phenotypic distances of two consecutive fragments, relative to the phenotypic distance from the empty line to the largest construct in the series, using independent measurements for the *broad*, *stripes*, *dimorphic*, *wing blade* and *spot* activities. To represent regulatory information, we used the absolute value of the change in the measure of activity, resulting in a similar depiction of repression and activation.

#### Abdominal transcriptome

##### RNA isolation

RNA isolation was performed on wild type flies using standard Trizol (Invitrogen) protocol from abdominal dorsal epithelium of pupae 72 hours after puparium formation. The Resulting RNA was further purified using “RNeasy mini Protocol” for RNA cleanup (Qiagen). RNA concentration was measured by spectrophotometer, and RNA quantity, quality and size distribution were checked on a Bioanalyser (Agilent Technologies).

##### Microarray experiment

In microarray experiments, each tissue was analyzed in three replicates consisting of 25 individuals each. RNA amplification was done according to Affymetrix “One cycle cDNA synthesis” and “Synthesis of biotin-labelled cRNA” protocols. Microarray hybridizations were conducted by Affymetrix facility.

Supplementary Text

Figs. S1 to S3

Tables S1 to S4

References (*40–43*)

**Fig. S1.**
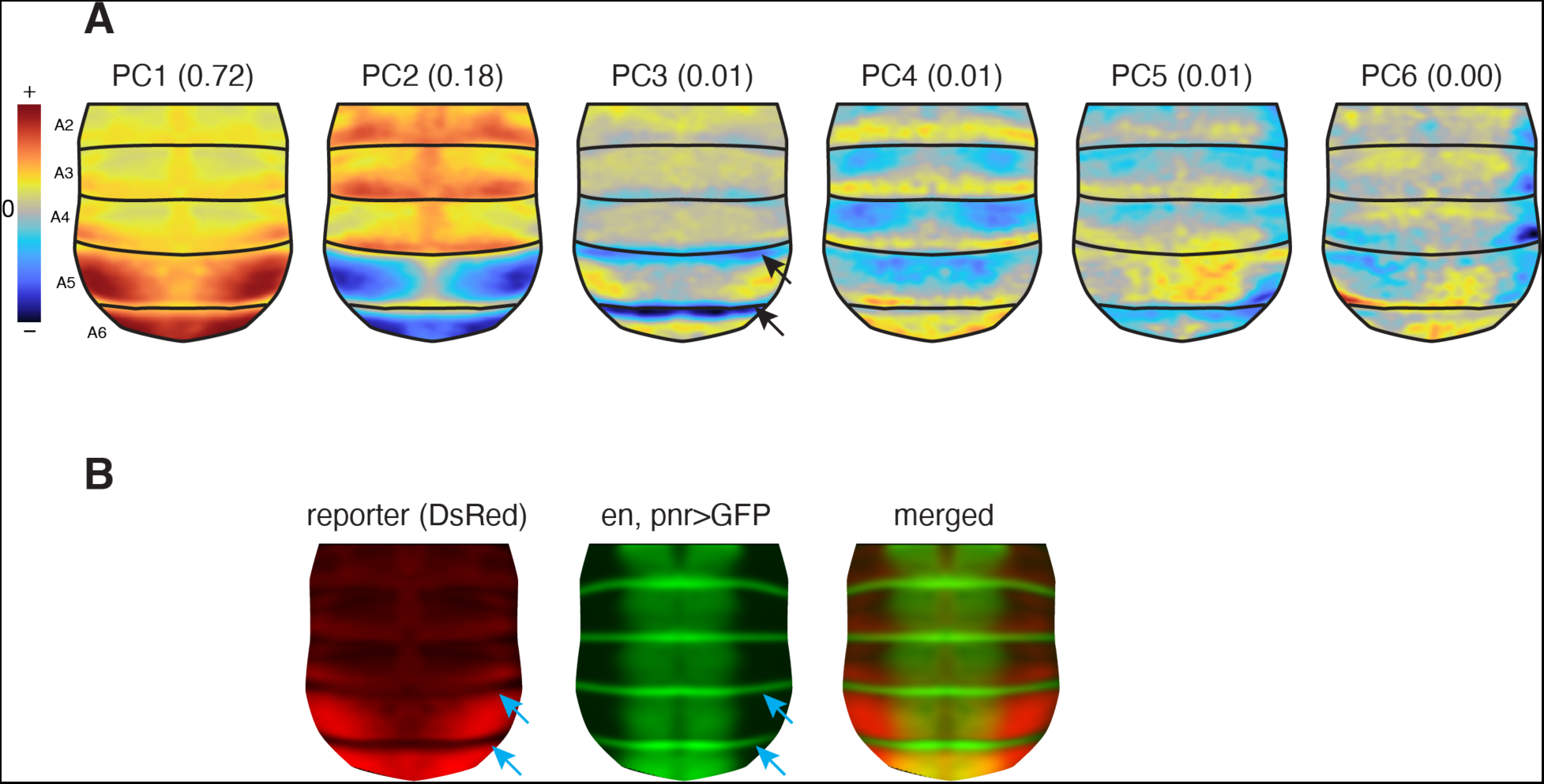
Directions in the principal component analysis of reporter lines. (**A**) Phenotypic directions of the first 6 principal components. PC1, PC2 and PC4 appear to capture biological information directly related to the regulatory dissection (see main text), PC3 captures mostly the gap between posterior segments (arrows) caused by their ventral bending and stretching, and PC5 and PC6 capture experimental noise (note the asymmetrical patterns). (**B**) The intersegmental gaps between segments A4-A5 and A5-A6 (blue arrows pointing to identical positions as in (A)) are visible on an average intensity projection of all reporter line images used in the PCA. For each reporter line image (DsRed), we also acquired a second image of fluorescent landmarks used for image registration and delineated by the combined expression of *engrailed*-Gal4 (*en*, segmental stripes) and *pannier*-Gal4 (*pnr*, longitudinal band), driving a UAS-GFP transgene. The merged image shows that the gaps captured by PC3 correspond to the posterior compartment of each segment defined by the expression of *engrailed*, an unpigmented domain of soft cuticle (*43*).

**Fig. S2.**
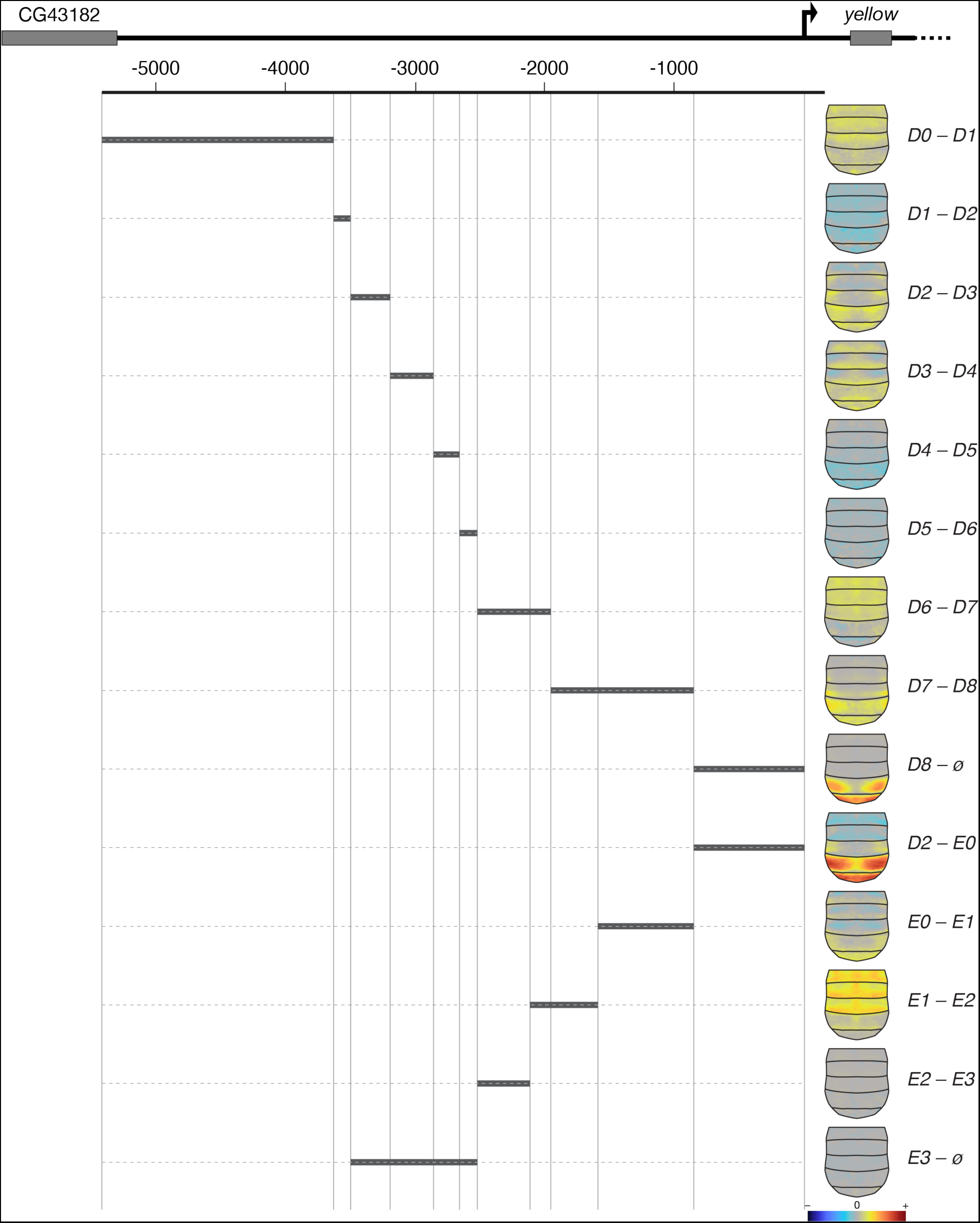
Subtraction of average phenotypes between consecutive lines highlights potential enhancer cores. Each horizontal gray segment represents the difference between two consecutive constructs. For instance, the difference *D0* – *D1* result in a distal-most segment. The abdomens on the right represent the calculated phenotypic difference between the average phenotypes of the corresponding lines, where differential fluorescent levels are indicated by a color map.

**Fig. S3.**
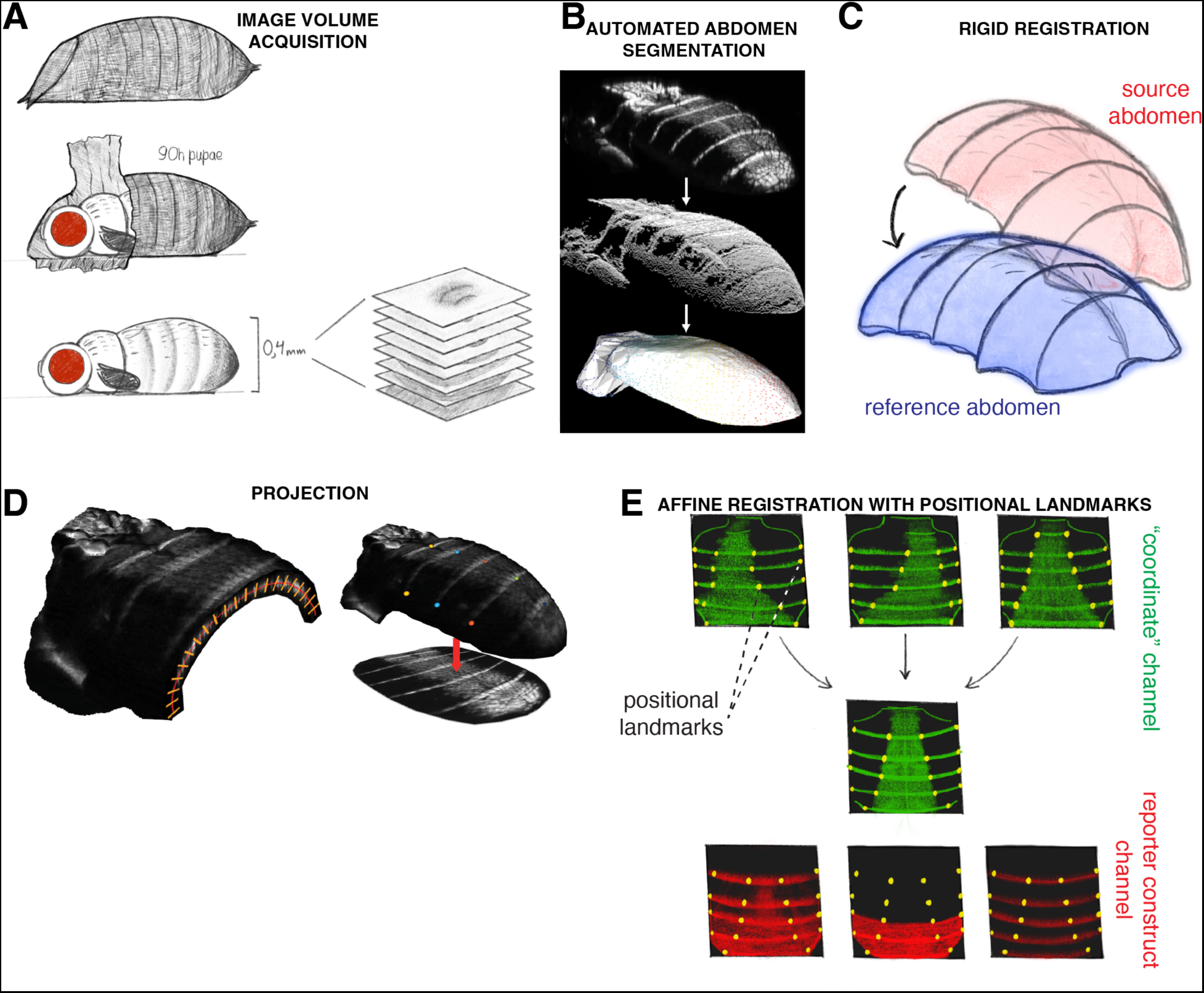
Pipeline for the registration, segmentation, and analysis of multi-channel 3D fluorescence microscopy images. (**A**) Acquisition of a 3D image volume of a *Drosophila* pupal abdomen (**B**) Segmentation of the abdomen from the 3D volume. (**C**) Rotation and rescaling of the source and the reference objects (rigid registration). (**D**) 2D projection of the 3D object using a modified sinusoidal projection. (**E**) Registration of the 2D images with user-placed positional landmarks (yellow dots).

**Table S1. Statistics for among-line expression comparisons.** Comparison of expression changes (fluorescence levels) between the reference line for each of series and the trimmed/randomized lines, and comparison between adjacent lines. P-values were obtained using the Kolmogorov-Smirnov test.

**Table S2. Wing vs. abdominal epidermis TF comparison.** The list of TFs expressed in pupal wings is the intersection of two published datasets: the micro-array dataset from the wings collected at 80 h after puparium formation (*36*) and RNA sequencing results from 72 h pupae (*37*). This list was compared to the TFs expressed in 72 h after puparium formation abdominal epidermis (previously unpublished micro-array dataset, see methods). TFs shared between the two tissues are highlighted (overlap wing/abdomen).

**Table S3. Primer sequences.** Primers for cloning of the *D8*, *Δ stripes* and *Δ broad* lines. Primers include additional sequence, creating overhangs for In-fusion cloning.

**Table S4. Sequences of the reporter constructs.** Sequences of the *D8*, *Δ stripes* and *Δ broad* lines. Randomized sequence fragments are underlined.

