## Supplementary material for "Entangled and non-modular enhancer sequences producing independent spatial activities": Table S4

**Table S4. Construct sequences.**

>D8

AAGCCAAGTCAATCGAAGCCCAGGTAATTCATATTTAGTGCTGTTCGCAAAGACCTGTCCCAGATACTCTGTTTATAGGTATAATTATTAAGTGCATATCAGGTTTATTTACATTTATATCGTATTATATTGGTAACTGCAGCAGATGCTGTGCTACAAATTTAGAATCATTTAAAACAAACATATTTGCCACAGAAAATGTGTGAAATAATTAAACTAAAAGCTTTGGATGAAGTAAAAAAGCCATAAAGCCTAAAATAATATTATGAATAATCAAAGAAAATCAGTAGATGGTAAAGTACTTCGTACCTACGTTGCATGGTATTCAATAAAGACTCGAAAATACTCTCACTCACTGTAAGTGAACCCAGTGTTTTGTAATTGCCTAGCACATAAATCAGCTGAATCCTAAACGTATCTGAAGGCCAGGAGTGTCGGAGAATTCGGTGTGCCAAAAACCAAAGACCAAAGACCATACCCTTTCAAAACCTTATGAAAAATGGCAAGCCCGGCGAAAGGTGTTGGCCGGTCCAGGGGATTCGGGGGCCCGTGATACTCGCACTTAATAAACATGCGTGAAAATCAATCAGCGAAGACAAAAGCCACGCACTAGAAGAAGCCAAAGTGTCCGAAGTGGCCGATCCACGGGTGACCATATAGACCATAAAGTCCGCATGGTGGACCACCACCCGAGCCACCGAAAGCAGCCGAATGGCCGAAACCCCGAAGTTGGCGCCTTCGTTTTCGCTTCCATTGGCCTGCCTTCGTCTTCGGAGAAAAAAACCTCATATAAAACGTGGCCGACATATTGAGTCCAACAGTCGTAAGCGCGCCACGGTCCACAGAA

>*Δ stripes*

AAGAGCCCAAGGTCGGTCGTTTAGCTTGGCCAAAACCTACCCATCCAGCTGGCACTTCCACCAACGGCACCAAGACACGAGCGGAAAATAAAAAGCCACACCACCCCACTTAGAACTCCGTTTAGCAGCAGTTGTTCAAACAGAAATTGGCTGGCTTCGGCCGGTTCAGCcTCAGTTGATAATTATTATAATATCTATGTTCTTGCCTATCGCTGGGCCCTAATTGGCCCAGACAAAGGCACCGTTTTTATGCATAACTGGAGGCTTACAATTTGGCcTTCGACACGCGGTCCTTCACATTGCCAAAAAAGAACGAGAACTCgGCAAGCCAATTACACTCGAAGAAGCAGCGGGGATCGTTTCGATGCCTAGCCCTGGGCCAGTTCAATCACTCCCGCCGATAATTAGCCGGCCTCTGCAATGGGAACTTTTCCAAGACGAGATCGATTCTTGGGAAAACACCACCTCAGTTTCCATTTCTGTTTTTTGGGTCCGGAAGTGGCATCGTGTCTTCCCAGAAGCGCCTCCAAATGGTGCCACCATTAGCCAGGGGGAAGCCGGGCGGCAGTCACTTAGCTGCTGCAATTTAAATACTTTTTAATTGATTACTAATTGCGGCGAGGCAAGTGCCAAAACAAGACGACGAGGACGACTTGGCTGTGACGTTTTCGATGCAACCCGACCGGGGACTGCCACTCTTTAGCCAGTTAATTGGCAGCAAAAGCGACAGTGGCAGCGGCAGCAACTGCTTTTCACCAGGAAATCAATAAACGCTCGTCCAGCGGCAAAAGTAATCGCAACACGCACATCTCAATTTCGGTGGCAGAAAAAAAACCCTCACCAGCTCAGTTCCCCGTGCCAAATTAACCAGAGCAAACATAGCCCAGTTTCTTCTCCTGCGGCATGTGAAAAGGCAAACAGTGCTGGCCGGAAAATACCCAGCAAAACACCTGAGTTCTAGTTGCGATTTTCGGAATTGGACTATAAAAGGCGGCCGTCGGGTAGCTTTCTTCACTCACAACCAGTCCAAAAGCATCTCCAACATGAAGTTCTTCCAGCAAATCGTCCTGGGTCTGATGCTCGTCCTGGCCATCATGGGTTCGCTGGCTAGCGCCAAGCCCCAGGAGGCCGAGGAACCGGAGGAGTCGCTGGTCGGGGACTCAGAGTCGGGACAGTCCGTGCCCGAGGACGCCCAGCAGGACTACCTCAACGTGGCGGACCTCACCACTGCCGCTCCTACATGGTGGTGGAACTAGAGCCCGGGAATTCTAGAGGATTTTAACAATCTTTTGTTTTTTTTTGCTAATCTAATGTACTATAATTGCAAAGAATTTACGGTGTTCCATATTCAATAAACCTATTTAAAGCTGAATACAATATTTACGTTAATAAATGTTCTTGATACGATAAATTTACTTAATTAAAATACATTACATTTCAGTTAAATATTTGTAAAATAAAATATATTTAAAAAAATATTTAATTCACTAGTTGTGGGAGTTCATTAGACTTATTATTTGTTTTTATTAAATTGTAATTTGTATCAAAAGTTTATTTTGCCAAACAGTGAATCTTAAAATATATATCAAGTTCATTGCACAAATTAACTTATAAATTGTCACCAAAAATTTAGAAATCAACCTATGTAAATAAATTTAACAAACCAATCATATCTTGAATTTAAATATATAAAAGAGGAGATAAACCATTTATAAAATGGTCTCACCTTTTTTTAGTTTATTTGATGCATGTTTTAATTTTGCTAAAATCATATTCTGATGTCTATTCATTTTGCCAGCCAGTAATCTAAAAAGTCGACCTATCACTCTCCCCCTCTTATATTTCGACCTATAAATACCCACCGCAAATGCCGCAACCAACCTAACCCACAGAGCCAGTTCGGGTTGTTTAATGGACAATTATCCTTAGTTCAGAAGGCGCCTGCCTTTATGCGTATTTCCCCGCTTGCCTGCGAATACGCCTAACGAAATTAATCGAGCCCGTAAACCCAGTTTCGGTAAGTGCTTCTTTATGAATATTTTCCATTTACTTTAATTGAAGGCTGCCAATTGTGGTGCCCGAGTGTTGTGACTGCTGGCCAATGAGGCGGTAATACGTTAAGTCGGAGCTGCGGAACGGGATGATGGACCAGTTGAGGCGAAGTACATCAATCTCATTTGCCCGCACTTATCGAACGGTTGCCTTGGGGAGATCGCTGCGATTGTTTATGAGTTATGTCCTGTGTCAGCGTAGATCGTTAATAACGCGAACGGTGGCCCACCCACAATAATGTGGGAGGCCACATACATGCTCACCATCGAACTCACGATTGCTGTGGTATACGTTTAGGGCGACCGAACGGAGTGTTAATGGTTATCCACCCGGCTGCCACATGATTCAGTGCGATGCGTACCCGAAGATATTCTCGGGAACCCAAGTAACTTATCGACATTTAATTGGCGGTTCCCCTTGCAATGCCTGGGTTATATAAACGACATATATTTATGTTGGGTCTTCTATACTCTGCGTAACAAGTAAATCCTTTCCCTCGGTGTCTGTTATCTAATTATTCCGTTTAAGGACGCAATTTTCTGAGCTAAAACTCGCTTATGGAGAGATCTAAATTTCCCCGCTTTTGGCTTGAATAAATTAATCGAATTCCCCGCTGGCTATTAAAACACACAAAAGGCGCTCTCGTCTGTTTCAATGTAAATTGCAAATTGCTCAATCCGCCTAATTGATGTGCGCCCATGCAATAGTTTTGTGCCAATCATTTTTAGTACACCCCTAACTGGTGTTTTCTACGCATAATATGTGCCATGGCTTAGGGCCTTTTGGTGGACTTACCAACTGAAGAAGACGATTGTGGGGGTGCGTTTGGCGCAGTGCGCGCCTGCGAGCAGGAAATCTCTTTCTCGGCCTGTCTGATTTTGGCCAAGACAAATAAATCCGGCTGGCAGATAGGCAGAGGGGACCCGGCGGTCAGGGCCGTGGACATTGAACTTGAAAACGCAGCCAGCGCCGAAAACATTGTATTCAACGAACGGCAAGTGCTGCGCGGCATGGGTGTCTCTGGCTAAGGTTACGGCGGTTGGGCAACAGGTTTTCCCCCGGCCAACACTGGGGGGAGAAAATAAAAAGGAAAATGTTCAGGCTGCCATAAGTGGGGAAAAAGGAAAACAAAACATGAAACACGGGCCGGGCAATGTCACTCGGCATTCGCTTGATTTTCCGCCTAACTCGCAGCGGTCCTGTGTGTAAATAATGTCTAATGTTGCATGCCGGTTGCATAATCGTGTGGCAATTATGCCAGAGAGATTCGCTTATTTATTTTTTACTTTCTGCCATGTTCCGCTGCCACCGTATTTCTTTTCGGCCACTTAGTGCGCTCCGCTTGATAATGATGTTTTGTTTTTCGCCGGGACAAACTCGTTTCGATTATTGGGAAAAGCGCGTATAAATCATCGCCGCCGAAGTCTGGCAAAACAGCAAATTGAAAACTGCAAGCTGAAAACTGAAAACTGAAAACTGTAACCCAAACAAACACAGCATCCCACACGACGAGGTGAAAATGAAAATAAATACGGACTGAGCGACTGAAAACGAGTCAATTCGATTCAAATTGCAGGTTCAACGGCTGCCGGCGATCGCATCATTAAGTGCGCCTTCGCTGGATACGCGGCTCTTATGCAACGAGCACACACAATTAATTAATAAGCGTCTGGTTGTTTCGGCCTGGCTTTTGCGGACCTGCCGATCGCAATAAATTAAGGCAGCATTAGTCGCAATTATGTGCCACATAGTTGGGCTGCTTACTTTTCTGTGGGTGAGCCGAGCGCAGAATGCGGCCAAGGGATCGAGTTAAACCGCTTTTCCGCAGGCCAAGAGTTTTTCGCATTTTGCATAAAATCGGCAACGCATAAGTGGCGAAGCATTGATGAAACTGCGGGAAAAGAAGTAAAAAATATTTAAAAATAAATATAAATTTATGGCAGAACTTAAGAAACTAATTTGAAATACTTCTTCTTAGGAAACTGTCCCTAGGAATATTTGTTTTCCCCAGCATTGCTCAATATTTCCTCCATCTTTTTGCTTATTGCCCAGACATTTTCCTTGGCCGAAGTGTAGCTGGTGGGTCTCCAGATTAATGCAAACCACTTCGTCAGCGGAGGTCGTAAACGTATCTTTGCCCATTTGGCTCGTTCATTATGCGTGTGGTATAGCTTTATTTTTGCCATTTTCCCTCTTTTTTGCACCAGCTGCAGTTGGGCCAAGAGAGTTATGCGAATCGGTGCGATTTTCGGGTTTTCGCACTCGCTTGCGGCCATGGCCATTAGAGCATTACCCGCTTAGGGCGCCCTAAAGTCCAGGTGGTCCCCAGGGACCACAAGAGTATTGCAACTTACGGCCAGCTGAGTGGAGTGCTGGAACGCACTTCTTAATTTCGGCGGTTATGTAACCTCGAGCTGAGTGTGCGATACATATGCCAAAATCACCTGCTCATAATTAGCGGAAACCAACTGTTTGGCCCTCGCCGGACTGTGAATCATCaGAGCTGCCCAATCGAAATCAAAGCCAAGTCAATCGAAGCCCAGGTAATTCATATTTAGTGCTGTTCGCAAAGACCTGTCCCAGATACTCTGTTTATAGGTATAATTATTAAGTGCATATCAGGTTTATTTACATTTATATCGTATTATATTGGTAACTGCAGCAGATGCTGTGCTACAAATTTAGAATCATTTAAAACAAACATATTTGCCACAGAAAATGTGTGAAATAATTAAACTAAAAGCTTTGGATGAAGTAAAAAAGCCATAAAGCCTAAAATAATATTATGAATAATCAAAGAAAATCAGTAGATGGTAAAGTACTTCGTACCTACGTTGCATGGTATTCAATAAAGACTCGAAAATACTCTCACTCACTGTAAGTGAACCCAGTGTTTTGTAATTGCCTAGCACATAAATCAGCTGAATCCTAAACGTATCTGAAGGCCAGGAGTGTCGGAGAATTCGGTGTGCCAAAAACCAAAGACCAAAGACCATACCCTTTCAAAACCTTATGAAAAATGGCAAGCCCGGCGAAAGGTGTTGGCCGGTCCAGGGGATTCGGGGGCCCGTGATACTCGCACTTAATAAACATGCGTGAAAATCAATCAGCGAAGACAAAAGCCACGCACTAGAAGAAGCCAAAGTGTCCGAAGTGGCCGATCCACGGGTGACCATATAGACCATAAAGTCCGCATGGTGGACCACCACCCGAGCCACCGAAAGCAGCCGAATGGCCGAAACCCCGAAGTTGGCGCCTTCGTTTTCGCTTCCATTGGCCTGCCTTCGTCTTCGGAGAAAAAAACCTCATATAAAACGTGGCCGACATATTGAGTCCAACAGTCGTAAGCGCGCCACGGTCCACAGAA

>*Δ broad*

AAGAGCCCAAGGTCGGTCGTTTAGCTTGGCCAAAACCTACCCATCCAGCTGGCACTTCCACCAACGGCACCAAGACACGAGCGGAAAATAAAAAGCCACACCACCCCACTTAGAACTCCGTTTAGCAGCAGTTGTTCAAACAGAAATTGGCTGGCTTCGGCCGGTTCAGCcTCAGTTGATAATTATTATAATATCTATGTTCTTGCCTATCGCTGGGCCCTAATTGGCCCAGACAAAGGCACCGTTTTTATGCATAACTGGAGGCTTACAATTTGGCcTTCGACACGCGGTCCTTCACATTGCCAAAAAAGAACGAGAACTCgGCAAGCCAATTACACTCGAAGAAGCAGCGGGGATCGTTTCGATGCCTAGCCCTGGGCCAGTTCAATCACTCCCGCCGATAATTAGCCGGCCTCTGCAATGGGAACTTTTCCAAGACGAGATCGATTCTTGGGAAAACACCACCTCAGTTTCCATTTCTGTTTTTTGGGTCCGGAAGTGGCATCGTGTCTTCCCAGAAGCGCCTCCAAATGGTGCCACCATTAGCCAGGGGGAAGCCGGGCGGCAGTCACTTAGCTGCTGCAATTTAAATACTTTTTAATTGATTACTAATTGCGGCGAGGCAAGTGCCAAAACAAGACGACGAGGACGACTTGGCTGTGACGTTTTCGATGCAACCCGACCGGGGACTGCCACTCTTTAGCCAGTTAATTGGCAGCAAAAGCGACAGTGGCAGCGGCAGCAACTGCTTTTCACCAGGAAATCAATAAACGCTCGTCCAGCGGCAAAAGTAATCGCAACACGCACATCTCAATTTCGGTGGCAGAAAAAAAACCCTCACCAGCTCAGTTCCCCGTGCCAAATTAACCAGAGCAAACATAGCCCAGTTTCTTCTCCTGCGGCATGTGAAAAGGCAAACAGTGCTGGCCGGAAAATACCCAGCAAAACACCTGAGTTCTAGTTGCGATTTTCGGAATTGGACTATAAAAGGCGGCCGTCGGGTAGCTTTCTTCACTCACAACCAGTCCAAAAGCATCTCCAACATGAAGTTCTTCCAGCAAATCGTCCTGGGTCTGATGCTCGTCCTGGCCATCATGGGTTCGCTGGCTAGCGCCAAGCCCCAGGAGGCCGAGGAACCGGAGGAGTCGCTGGTCGGGGACTCAGAGTCGGGACAGTCCGTGCCCGAGGACGCCCAGCAGGACTACCTCAACGTGGCGGACCTCACCACTGCCGCTCCTACATGGTGGTGGAACTAGAGCCCGGGAATTCTAGAGGATTTTAACAATCTTTTGTTTTTTTTTGCTAATCTAATGTACTATAATTGCAAAGAATTTACGGTGTTCCATATTCAATAAACCTATTTAAAGCTGAATACAATATTTACGTTAATAAATGTTCTTGATACGATAAATTTACTTAATTAAAATACATTACATTTCAGTTAAATATTTGTAAAATAAAATATATTTAAAAAAATATTTAATTCACTAGTTGTGGGAGTTCATTAGACTTATTATTTGTTTTTATTAAATTGTAATTTGTATCAAAAGTTTATTTTGCCAAACAGTGAATCTTAAAATATATATCAAGTTCATTGCACAAATTAACTTATAAATTGTCACCAAAAATTTAGAAATCAACCTATGTAAATAAATTTAACAAACCAATCATATCTTGAATTTAAATATATAAAAGAGGAGATAAACCATTTATAAAATGGTCTCACCTTTTTTTAGTTTATTTGATGCATGTTTTAATTTTGCTAAAATCATATTCTGATGTCTATTCATTTTGCCAGCCAGTAATCTAAAAAGTCGACCTATCACTCTCCCCCTCTTATATTTCGACCTATAAATACCCACCGCAAATGCCGCAACCAACCTAACCCACAGAGCCAGTTCGGGTTGTTTAATGGACAATTATCCTTAGTTCAGAAGGCGCCTGCCTTTATGCGTATTTCCCCGCTTGCCTGCGAATACGCCTAACGAAATTAATCGAGCCCGTAAACCCAGTTTCGGTAAGTGCTTCTTTATGAATATTTTCCATTTACTTTAATTGAAGGCTGCCAATTGTGGTGCCCGAGTGTTGTGACTGCTGGCCAATGAGGCGGTAATACGTTAAGTCGGAGCTGCGGAACGGGATGATGGACCAGTTGAGGCGAAGTACATCAATCTCATTTGCCCGCACTTATCGAACGGTTGCCTTGGGGAGATCGCTGCGATTGTTTATCGATAATCGCCCGATTACCGCGCTGAGCGGTCTTAAAGACCCATAAGAAATGCGGCGATGGCGGCTTTAGATAAGTAAGTCGTCGGGGCGCTCATAAATTTCGAGCGCGATCCACGGATTTATGCACTCGCTGGAAAAGCTATTACCATTAGGCTTTTCGCGACCACGGATTTTTCCGCTTGCCTGAGACAAGTGCAGCGCGGCAGTTGCAGGCAAATTATGTGTGAGGCAATGCCGCGGGCATGTCTACACCGAAATCAAATTACGGCAACCTCTATTCACTTATTTGCTTAGTTTTTTGCGCAGTGAGCGGCCAGCGCCTTGTATTGGCATCTAATTATTCCGTTTAAGGACGCAATTTTCTGAGCTAAAACTCGCTTATGGAGAGATCTAAATTTCCCCGCTTTTGGCTTGAATAAATTAATCGAATTCCCCGCTGGCTATTAAAACACACAAAAGGCGCTCTCGTCTGTTTCAATGTAAATTGCAAATTGCTCAATCCGCCTAATTGATGTGCGCCCATGCAATAGTTTTGTGCCAATCATTTTTAGTACACCCCTAACTGGTGTTTTCTACGCATAATATGTGCCATGGCTTAGGGCCTTTTGGTGGACTTACCAACTGAAGAAGACGATTGTGGGGGTGCGTTTGGCGCAGTGCGCGCCTGCGAGCAGGAAATCTCTTTCTCGGCCTGTCTGATTTTGGCCAAGACAAATAAATCCGGCTGGCAGATAGGCAGAGGGGACCCGGCGGTCAGGGCCGTGGACATTGAACTTGAAAACGCAGCCAGCGCCGAAAACATTGTATTCAACGAACGGCAAGTGCTGCGCGGCATGGGTGTCTCTGGCTAAGGTTACGGCGGTTGGGCAACAGGTTTTCCCCCGGCCAACACTGGGGGGAGAAAATAAAAAGGAAAATGTTCAGGCTGCCATAAGTGGGGAAAAAGGAAAACAAAACATGAAACACGGGCCGGGCAATGTCACTCGGCATTCGCTTGATTTTCCGCCTAACTCGCAGCGGTCCTGTGTGTAAATAATGTCTAATGTTGCATGCCGGTTGCATAATCGTGTGGCAATTATGTTGATGTATCCTGATTAAGCTAGGTCTCTTTGAACTTGTGCAGCTCCACGGGATAGCCGAACGTTTCGGAGTTTGTGTGTCTTTCTTCCATATGCTTCGTGTAATTACATTTATTCCACAAACAATAAAATAGAGGGGACCTGTCTAAAGAACAACACATGGCAAAGTGGGAATACAACCAGAAAAGTGGTCCAATAAACAAAGAACGTGAATCACTCAGGAATGAGAACCATCGTGAGCCTTCAGCAACAATTACCCATGGCATCTAAATGGCGAGTACTTTACAACGCCTGACAAAGATAGCTTACGAATCATGTGACGCGAGTATCAATAATTTTGTATGAGTCTCACCCAGATTCTGATCCGCCGTTAAGCTCACCCGTTAGGCAAACTCTTTGGCATCGATGGTAGTTAGCTCCATGTAAACAATTCTTACTAGAGGTAGGCCCAGCGTGCGCGCGCTTACCTATTGAGGGTTTGATCGCCCTTTAGTAGAGTCGGGGTCCGGCTTCAGGTATCGAATCGAGCGCAGAATGCGGCCAAGGGATCGAGTTAAACCGCTTTTCCGCAGGCCAAGAGTTTTTCGCATTTTGCATAAAATCGGCAACGCATAAGTGGCGAAGCATTGATGAAACTGCGGGAAAAGAAGTAAAAAATATTTAAAAATAAATATAAATTTATGGCAGAACTTAAGAAACTAATTTGAAATACTTCTTCTTAGGAAACTGTCCCTAGGAATATTTGTTTTCCCCAGCATTGCTCAATATTTCCTCCATCTTTTTGCTTATTGCCCAGACATTTTCCTTGGCCGAAGTGTAGCTGGTGGGTCTCCAGATTAATGCAAACCACTTCGTCAGCGGAGGTCGTAAACGTATCTTTGCCCATTTGGCTCGTTCATTATGCGTGTGGTATAGCTTTATTTTTGCCATTTTCCCTCTTTTTTGCACCAGCTGCAGTTGGGCCAAGAGAGTTATGCGAATCGGTGCGATTTTCGGGTTTTCGCACTCGCTTGCGGCCATGGCCATTAGAGCATTACCCGCTTAGGGCGCCCTAAAGTCCAGGTGGTCCCCAGGGACCACAAGAGTATTGCAACTTACGGCCAGCTGAGTGGAGTGCTGGAACGCACTTCTTAATTTCGGCGGTTATGTAACCTCGAGCTGAGTGTGCGATACATATGCCAAAATCACCTGCTCATAATTAGCGGAAACCAACTGTTTGGCCCTCGCCGGACTGTGAATCATCaGAGCTGCCCAATCGAAATCAAAGCCAAGTCAATCGAAGCCCAGGTAATTCATATTTAGTGCTGTTCGCAAAGACCTGTCCCAGATACTCTGTTTATAGGTATAATTATTAAGTGCATATCAGGTTTATTTACATTTATATCGTATTATATTGGTAACTGCAGCAGATGCTGTGCTACAAATTTAGAATCATTTAAAACAAACATATTTGCCACAGAAAATGTGTGAAATAATTAAACTAAAAGCTTTGGATGAAGTAAAAAAGCCATAAAGCCTAAAATAATATTATGAATAATCAAAGAAAATCAGTAGATGGTAAAGTACTTCGTACCTACGTTGCATGGTATTCAATAAAGACTCGAAAATACTCTCACTCACTGTAAGTGAACCCAGTGTTTTGTAATTGCCTAGCACATAAATCAGCTGAATCCTAAACGTATCTGAAGGCCAGGAGTGTCGGAGAATTCGGTGTGCCAAAAACCAAAGACCAAAGACCATACCCTTTCAAAACCTTATGAAAAATGGCAAGCCCGGCGAAAGGTGTTGGCCGGTCCAGGGGATTCGGGGGCCCGTGATACTCGCACTTAATAAACATGCGTGAAAATCAATCAGCGAAGACAAAAGCCACGCACTAGAAGAAGCCAAAGTGTCCGAAGTGGCCGATCCACGGGTGACCATATAGACCATAAAGTCCGCATGGTGGACCACCACCCGAGCCACCGAAAGCAGCCGAATGGCCGAAACCCCGAAGTTGGCGCCTTCGTTTTCGCTTCCATTGGCCTGCCTTCGTCTTCGGAGAAAAAAACCTCATATAAAACGTGGCCGACATATTGAGTCCAACAGTCGTAAGCGCGCCACGGTCCACAGAA
